# Similarities and dissimilarities between psychiatric cluster disorders

**DOI:** 10.1101/2020.08.06.239566

**Authors:** Marissa A. Smail, Xiaojun Wu, Nicholas D. Henkel, Hunter M. Eby, James P. Herman, Robert E. McCullumsmith, Rammohan Shukla

**Author notes:** To whom the correspondence should be addressed: Rammohan Shukla, 3000 Arlington Avenue, room 182, Toledo, Ohio 43614, USA.

## Abstract

The common molecular mechanisms underlying psychiatric disorders are not well understood. Prior attempts to assess the pathological mechanisms responsible for psychiatric disorders have been limited by biased selection of comparable disorders, datasets as well as challenges associated with data normalization. However, publicly available databases offer a unique opportunity to expand such investigations both in terms of the number and types of diseases. Here, we used DisGeNET, a database of over 24,000 gene-disease associations to investigate the similarities and dissimilarities associated with enrichment of pathways, cell-types, drug targets, and human chromosomes within an unbiased cluster of psychiatric disorders. We show that cognition and neurotransmission related pathways are involved across all disorders, whereas those associated with immune system and signal-response coupling (cell-surface receptors, signal-transduction, gene-expression, and metabolic process) are associated with few disorders of the cluster. The drug-target based enrichment confirms the involvement of neurotransmission related changes across these disorders. At cell-type level, dendrite targeting interneurons, across all layers, are most involved across all disorders. Finally, using a clustering-based similarity index, we showed that the similarity between the disorders are influenced most at chromosomal level and to some extent at cellular level. Collectively, the results provide a comprehensive comparison of many psychiatric diseases in an unbiased manner and expand our understanding of the cellular and molecular pathologies associated with similar and comorbid psychiatric disorders.

## INTRODUCTION

Psychiatry encompasses a vast number of disorders and comorbidity. While these disorders have their own unique traits, common molecular mechanisms may be involved in their underlying pathology^1^. Identifying such common elements would enhance our understanding of numerous disorders simultaneously and identify common therapeutics against them.

Prior efforts to identify such common mechanisms were informative yet incomplete. Most studies^2,3^, in an attempt to make the data more manageable, compared the transcriptomic profiles of a few diseases at a time, limiting their ability to reveal patterns across a wide range of conditions and comorbidity. The diseases included in these studies were selected based on disease-severity known associations or data/cohort availability^4^, thus preventing the exploration of novel relationships or study of known associations (e.g. depression and epilepsy^5^). Furthermore, normalization often presents a challenge in these studies^6,7^, and efforts to unionize diverse datasets can compromise the results and limit the conclusions drawn from them.

Another approach to this comprehensive analysis is to draw on publicly available gene-disease databases. These resources have greatly expanded in number and detail in the past several years; thus, enhancing the number of diseases that can be compared, simultaneously. One such resource is DisGeNET^8,9^, a knowledge management platform cataloging genes associated with several human diseases (Mendelian, complex and environmental). Utilizing over 20 diverse resources, of human, animal, and computational data, DisGeNET identifies gene-disease associations (GDAs) and generates a list of genes associated with each disease. Presently, DisGeNET has curated disease-associated gene-sets for 24,166 diseases, featuring 628,685 GDAs across 17,549 genes^8^. As these gene lists are not reliant upon expression profile, it precludes the limitations that have compromised prior efforts to compare psychiatric disorders. Furthermore, the sheer number of diseases with GDAs makes this an ideal platform to compare several related and diverse diseases in an unbiased manner.

Here, we use curated and evidence supported, clinically relevant disease-associated gene-sets from DisGeNET; to identify unbiased clusters of similar diseases; then, with special emphasis on psychiatric disorders, we performed series of in silico analyses dissecting four different levels of biological complexity –pathways, cell-types, drug-targets, and chromosomes— to identify mechanisms that are commonly dysregulated in a cluster of psychiatric disorders. This top down approach is critical to identifying highly related disorders in an unbiased manner, revealing common and unique mechanisms underlying disease pathology.

## METHODS

### Disease-Disease Similarity

Curated disease-associated gene-sets were downloaded from DisGeNET. In order to avoid size related bias and improve the specificity as well as the interpretability of pathway profiles^10–12^, we restricted our analysis to 763 diseases with gene-set size between 10 to 500. Pairwise-similarity between disease-associated gene-sets was calculated using Jaccard similarity-index^13^ implemented by gene-overlap package in R.

### Filtering for Psychiatry Cluster

A three-step top down approach was used to filter for a psychiatric cluster. First, using principal component analysis (PCA) (FactoMineR package in R) of scaled Jaccard similarity-matrix, global clusters of disease were identified. Second, focusing on the PCA cluster enriched with psychiatric disorders, a Euclidean distance-based dendrogram was generated for disorders in this cluster. Finally, using the cuttree function, dendrogram clusters most enriched with psychiatric disorders were used for all further analysis.

### Gene-Ontology Analysis

Pathways affected in different diseases of the psychiatric cluster were determined using hypergeometric overlap analysis (HGA) with a background of 21,196 genes (default, gene-overlap package in R). The significant overlap (q-value<0.05) of disease-associated gene-sets were tested against gene-ontology (GO) pathways associated with Biological-Process (GOBP), Molecular-Function (GOMF), and Cellular-Component (GOCC). Updated list of GO-pathways were obtained from the Bader-lab (http://download.baderlab.org/EM_Genesets/). To compare the effect of pathways across different diseases, the -log10(q-value) was used to generate the heatmap. To better identify the character of biological changes, in the overlap results, a focused analysis of forty a-priori functional themes was performed. As describe in our previous study^14^, the pathways were filtered based on the parent-child association between GO-terms in our list of significant pathways (child-pathways) and hand-picked parent-pathways representing the a-priori theme from the GO-database (GOdb package in R).

### Density-Index

To quantitatively summarize how common (close to 1) or unique (close to 0) a theme is across different subgroups of psychiatric-disorders; we devised a density-index. For a given ***r* ⨯ *c*** matrix of -log10(q-value), a density-score is obtained as:

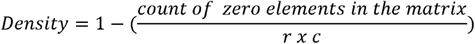

Where ***r*** and ***c*** represent the number of pathways in a theme and number of disorders in a psychiatric subgroup, respectively. A zero element represents a non-significant disease-pathway relationship. For density associated with individual pathways, cell-types, drugs and chromosomes a vector representing the number of disorders (=36) in the psychiatric-cluster was used. In such an instance, the numerator of the above density formula represents count of zero elements in a vector. Whereas ***r*** and ***c*** in the denominator are constant holding a fixed value of 1 and 36, respectively.

### Cell-Type and Drug-Target Enrichment Analysis

Enrichment of neuronal cell-types and drug induced molecular signatures in the disease-associated gene-sets was calculated using HGA. Human cell-specific markers from two different studies were used^15,16^. Drug specific gene-markers were downloaded from the Enricher library of gene-sets. In order to understand the druggable-mechanism and targets involved, gene-markers of drugs with known modes of action (MOA) and targets were used.

### Chromosome overrepresentation analysis

To access the chromosomal enrichment of each gene-set in the psychiatry cluster we used Fisher’s exact test (a base-function in R). A non-redundant list of genes within each chromosome was downloaded from Hugo Gene Nomenclature Committee^17^ and used as background.

### Rand-index

To compare the clustering of disorders observed in psychiatric-cluster with the clustering of same disorders using pathway, cell-type, drug-target, and chromosome enrichment profiles, we used Rand-index (fossil package in R). The cluster labels generated by each cluster were used as input for the comparison. Significance was generated by two tailed randomization-test using 1000 resampling permutations of psychiatry cluster labels as reference.

## RESULTS

### Disease profiles fall into three distinct clusters

To look for molecularly similar diseases, we used 763 disease-associated gene-sets from DisGeNET and calculated a matrix of pairwise-similarity (Table S1, methods). A PCA over this matrix segregated the disease profiles into three distinct clusters (Fig 1A, Table S1). Cluster-1 containing 422 disease profiles was positioned at the center of the plot, suggesting a low correlation between diseases in this cluster. Cluster-2 and Cluster-3 were orthogonal to each other and each contained a distinct subset of highly correlated diseases. Cluster-2 contained 192 profiles and was primarily composed of diseases related to psychiatric disorders, inflammation, metabolism, and neurodegeneration. Cluster-3 contained 149 profiles and was primarily composed of various types of cancer. Overall, the separation of disease profiles into 3 clusters suggesting that converging mechanisms are involved in presentation of diseases within each cluster. Given our group’s focus on psychiatric disorders, the remainder of this paper will focus specifically on Cluster-2.

**Figure 1:**
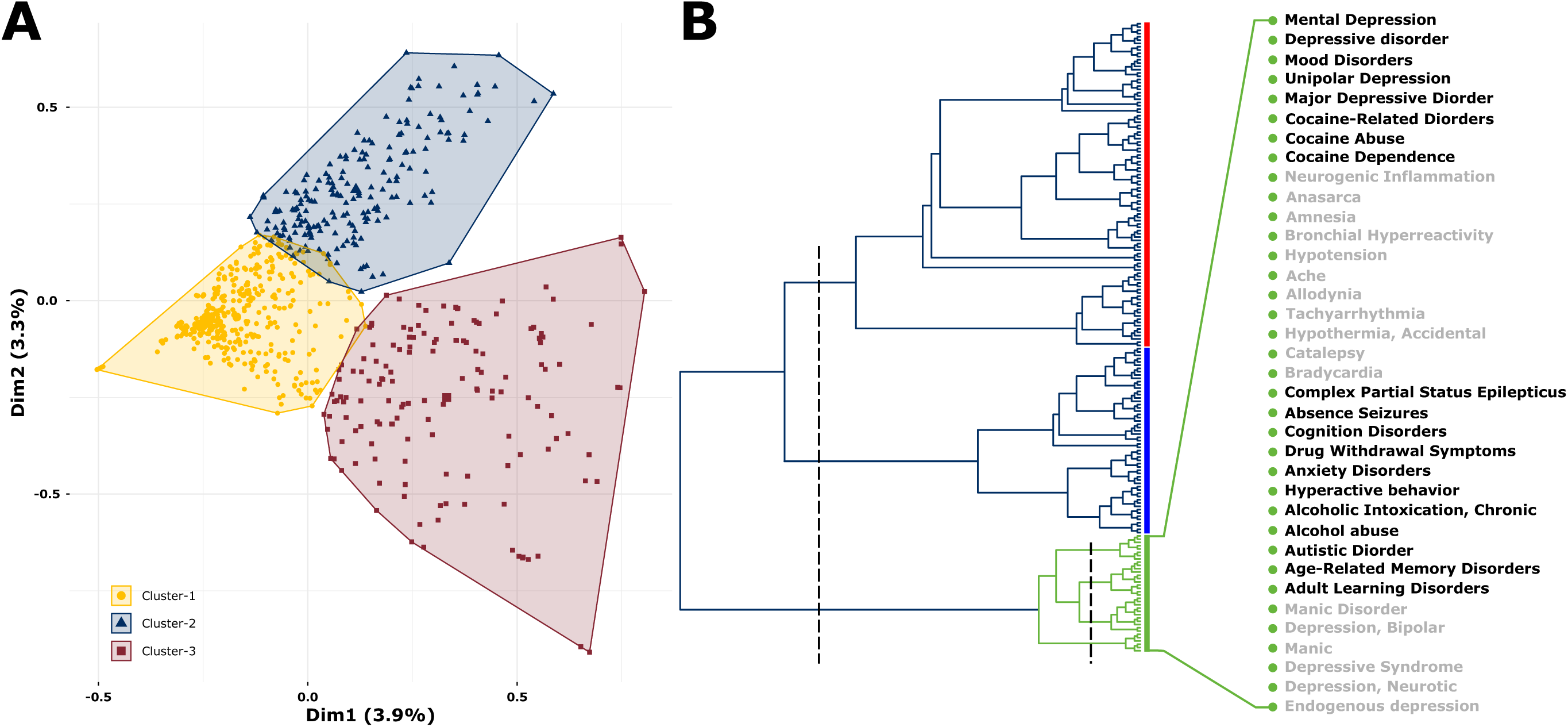
Disease-associated gene-sets fall into three distinct clusters. **(A)** A PCA based clustering of 763 curated disorders. Note that cluster-1 falls at the center of the plot indicating a low internal correlation between the disorders in this cluster. **(B)** Hierarchical clustering of Cluster-2 disorders. An initial cutting of three (left dashed line) reveals three branches shown in red, blue and green. The green branch (aka the psychiatric cluster) was considered for all subsequent analysis. Further cutting the green branch (right dashed line) reveals four subgroups composed of 36 disorders labelled on the right. See Figure S1 for further details.

### Hierarchical clustering reveals distinct subgroup of psychiatric disorders

Hierarchical clustering of Cluster-2 disease profiles revealed three distinct branches, (Fig 1B, Fig S1). Branch-1 (Fig 1B, red) was the largest and contained a range of diseases primarily related to metabolism, organ-failure, vascular-disorders, mood, and neurodegenerative-disorders. Subgroups within this cluster recapitulate known associations between diseases. For example, diabetes-related disorders largely cluster together, as do vascular- and neurodegeneration-related disorders. Branch-2 (Fig 1B, blue) was primarily composed of immune-system disorders. These included autoimmune diseases, multiple-sclerosis, Crohn’s disease, dermatitis, lung-inflammation, allergic-reaction, injury, and fever. Branch-3 (Fig 1B, green) was primarily composed of neuropsychiatric disorders, including depression, addiction, bipolar, anxiety, and learning-disorders. It also contained epilepsy, cardiovascular disorders, and pain, which are comorbid with psychiatric disorders^18–21^. Overall, these Cluster-2 branches show that shared mechanisms in branches are stronger than the original clusters, and the known mutual similarity of branch-revealed diseases gives confidence to selectively investigate new connections between disorders and their underlying mechanisms.

### Cognition is the most affected biological process across all psychiatric cluster disorders

The high concentration of psychiatric disorders in Branch-3 (Fig 1B green, henceforth referred to as the “psychiatric cluster”) makes it the ideal place to investigate underlying mechanisms across psychiatric illnesses; thus, it remained the focus for subsequent analyses. This branch splits into four distinct subgroups (Fig 1B green, expansion).

Gene-ontology (GO) analysis of gene-sets associated with diseases in psychiatric cluster revealed over 3000 GO-pathways (q-value<0.05, Table S3). To better understand the disease process, the pathways were organized into 40 themes representing different levels of cellular and biological complexity (Fig 2A, left labels). To facilitate a theme-centric quantitative comparison, we calculated a density-index for each theme (Fig 2 right, methods). A density close to 1 indicates themes common across diseases, whereas a density close to 0 indicates themes unique to a few diseases. Across the entire psychiatric cluster, *cognition* shows the highest density (density _≈_ 0.7). Other high-density (density _≈_ 0.5) themes include *neurotransmission*, largely driven by *catecholamine* and *serotonin*, and signaling pathways driven by *G-protein-coupled-receptors*. Medium-density (density > 0.25) themes include other neurotransmitters, including *glutamate, GABA*, and *norepinephrine*; ion balance, driven by *transmembrane-transport*; *post-synaptic events* and *inflammatory-response*. Low-density (density < 0.25) themes include those related to the *immune-system, metabolism, cell-surface receptor-signaling, intracellular-signal-transduction* and *oxidative-stress*.

**Figure 2:**
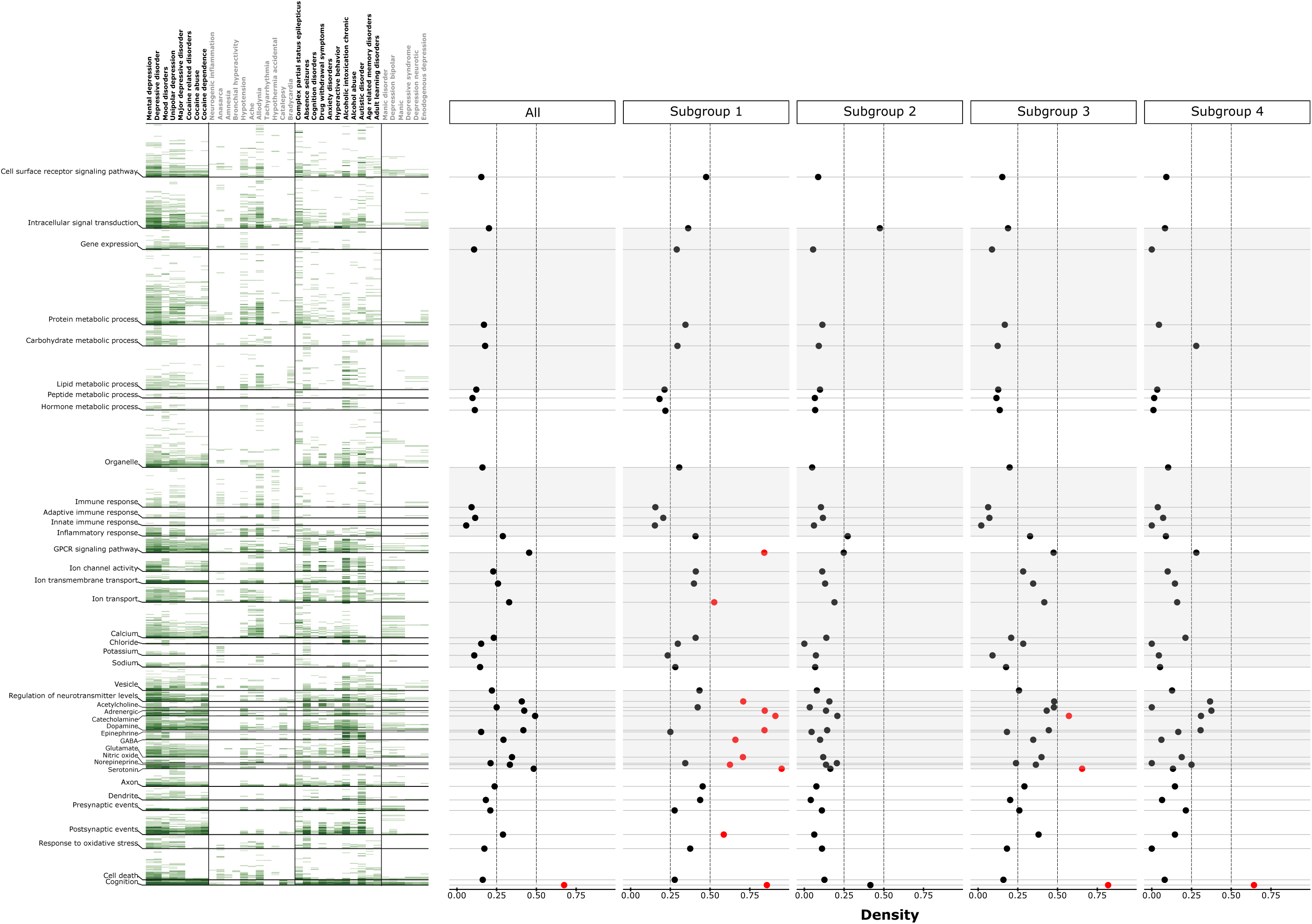
Pathway analysis of psychiatric cluster disorders. **Left:** Heatmap of significant (q<0.05) and theme (left labels) filtered pathways associated with all psychiatric cluster disorders **(top labels)**. The color-intensity (light to dark green) in the heatmap is proportional to -log10(q-value) **Right:** Densities of each theme across all disorders and individual subgroups. The red dots represent the highest density themes crossing the threshold (arbitrary) of 0.5. Note the highest density of cognition across all disorders.

While *cognition* is the most consistently affected process, comparing themes across subgroups reveals differences between the mechanisms underlying this phenotype. Subgroup-1, containing Major-depressive-disorder (MDD) and cocaine-addiction, exhibits the highest density indices across all the subgroups for themes related to *adrenergic, catecholamine, dopamine, GABA, glutamate, norepinephrine* and *serotonin*-related neurotransmission, possibly involving the slow *G-protein-coupled-receptor-signaling*, which too shows a high-density index. Interestingly, as compared to increased neurotransmission, subgroup-1 showed reduced *presynaptic* events relative to *postsynaptic* events, suggesting a reduced input functionality.

Subgroup-2, containing cardiovascular conditions and pain, features the lowest density of *cognition* and exhibits much lower density-indices across all theme-subgroup comparisons, suggesting subtle changes governed by unique mechanisms. Aside from cognition, the highest density themes in this subgroup are *G-protein-coupled-receptor-signaling, ion-channel* activity, *inflammatory-response*, and neurotransmission related to *nitric-oxide* and *catecholamine*. As indicated by the low theme-density correlations (Fig S2), this subgroup differs most from the other subgroups likely due to more relative contribution of immune system dysfunction than alterations to neurotransmission.

Subgroup-3, containing epilepsy, anxiety, alcohol-abuse, and ADHD, exhibits a density pattern similar to Subgroup-1 and both subgroups showed highest theme-density correlation (Fig S2). Density associated with cognition was particularly similar in both subgroups (_≈_ 0.8); however, other themes, though similar, showed low density. Besides *cognition*, the highest density themes are related to neurotransmission, largely driven by *serotonin* and *catecholamine*. Though not similar in density, *inflammatory-response, postsynaptic-events*, and ion balance showed changes similar to Subgroup-1. Overall, the similarity between the two subgroups indicates that alcohol and cocaine abuse have similar downstream effects and mechanisms widely shared with depression, anxiety, and epilepsy.

Subgroup-4, containing bipolar-disorder, mostly exhibits medium-density theme indices which showed similar correlations across subgroups (Fig S2). Neurotransmission associated with *adrenergic, catecholamine*, and *dopamine* was affected the most. Interestingly, density of *carbohydrate-metabolic-process* is uniquely high in this group, suggesting that metabolic processes may play a larger role in bipolar-disorders over other psychiatric disorders. Indeed, there are evidences for dysregulated metabolic processes in manic states^22,23^. Overall, the unsupervised-clustering of the highly comorbid psychiatric disorders suggests that neurotransmission, mostly associated with monoamines and governed by *G-couple-protein-receptors*, is the key shared process across psychiatric disease. Whereas pathways associated with *cell-surface-receptors, signal-transductions*, and *metabolic-process* appear to be more unique across a few disorders.

### Druggable mechanisms confirms the involvement of neurotransmission across psychiatric cluster disorders

Therapeutic or disease-inducing drugs with known MOA can expand our understanding of the underlying disease pathology^24^. By comparing the psychiatric disease-associated gene-sets with those of known drugs from the connectivity map, a database cataloging transcriptomic responses of several cell-lines against known drugs, we identified 132 relevant drugs, belonging to 64 different MOAs (Table S3).

The most frequent MOAs across the entire psychiatric cluster involved *dopamine-receptors* (15/64), *adrenergic-receptors* (14/64), *glucocorticoid-receptors* (8/64), and *ATPases-activity* (6/64). Interestingly, these MOAs remained most frequent across each subgroup, considered individually (Fig S3). Within each subgroup, Subgroup-1 showed the most with 44 MOAs, whereas Subgroup-4 showed the least with 13 MOAs. Subgroup-3 and Subgroup-4 showed 20 and 24 different MOAs, respectively. Looking at specific drugs, those with the strongest density were helveticoside (targeting *ATPases*), thioridazine (targeting *dopamine-receptors*), clioquinol (targeting *opioid-receptors*), and anisomycin (targeting *DNA-synthesis*). Overall, while the drug target analysis confirms the pathway analysis findings that most psychiatric disorders feature dysregulated neurotransmission, the diverse MOAs points towards the heterogeneous nature of psychiatric-disease origins.

### Interneurons are most affected cell-types across psychiatric cluster disorders

Alterations in various layer-specific subtypes of neurons and glia is an important factor driving disease mechanisms and could differentially contribute to pathology^25,26^. Using human-specific markers from two independent studies, with and without layer specificity, taken from middle temporal gyrus^16^ and anterior cingulate cortex^15^, respectively, we assessed the enrichment of neuronal and non-neuronal cell-types in the psychiatric cluster (Fig 3). Based on cell-specific markers (Fig 3, top), Subgroup-1 diseases were enriched in somatostatin (SST) and corticotrophin-releasing-hormone (CRH) positive interneurons in a non-overlapping manner. Mental-depression and cocaine-addiction were enriched in SST positive interneurons, whereas unipolar-depression and major-depression were enriched in CRH positive interneurons. Subgroup-2 diseases were enriched with more diverse cell-types and consistent with their role in inflammatory diseases, were also found enriched in neuroglia and oligodendrocytes. However, CRH and vasoactive intestinal polypeptide (VIP) co-expressing CRH neurons were most abundant in this cluster. Subgroup-3, consistent with the high theme-centric correlation with Subgroup-1, was also enriched in SST and CRH positive interneurons. In Subgroup-4, endogenous, neurotic, and syndromic form of depression, similar to mental depression observed in Subgroup-1, were enriched in SST interneurons, whereas those with bipolar-disorder were enriched in Parvalbumin (PV) -positive and CRH positive interneurons.

**Figure 3:**
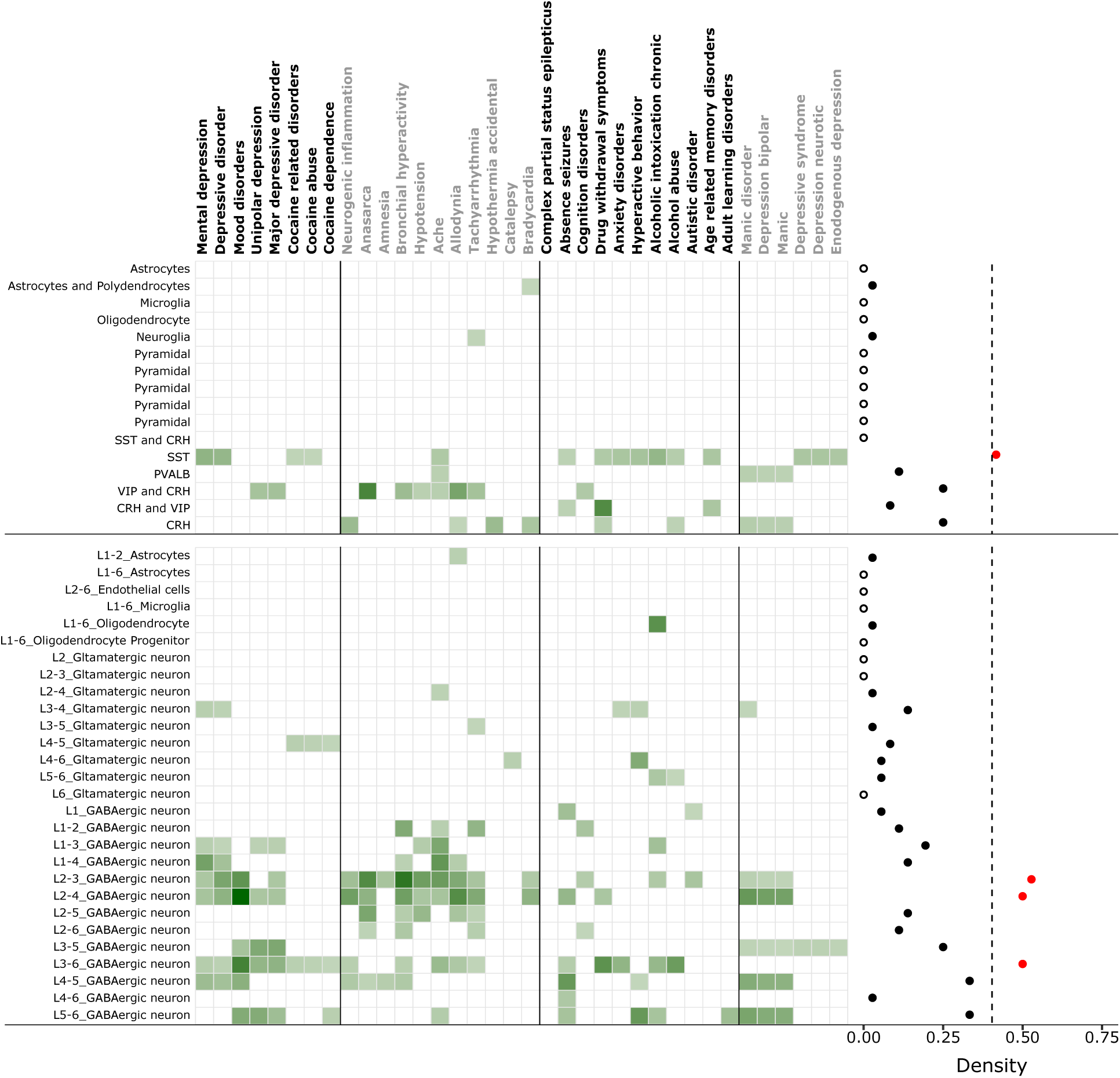
Cell-type analysis of psychiatric cluster disorders. Enrichment of different cell-types (top-left labels) and layer specific cell-types (bottom-left labels) markers across psychiatric cluster disorders (top labels). **Right:** Density of each cell-type (top) and layer-specific cell-types (bottom) across all disorders. The filled red dots represent high density cell-types whereas hollow dot represents a cell-type with zero density. Note the highest density of SST positive interneurons and layer 2/3 specific interneurons across all disorders. The color-intensity (light to dark green) in the heatmap is proportional to -log10(q-value).

Consistent with cell-specific markers, layer-specific markers, also showed excessive enrichment in interneurons, mostly distributed across all layers (Fig 3, bottom). However, based on cell-specific density-index calculated across all the disorders in psychiatric cluster, GABAergic interneurons of layer-2/3 show the highest enrichment (density = 0.52). Also consistent with cell-specific markers, associations with other cell-types were weaker and less consistent. Within glutamatergic neurons, those found in layer 3/4 showed the highest density of enrichment. Astrocytes and oligodendrocytes were involved with very few diseases with a density index of ∼0.02. Overall, holding true for markers from two independent studies, interneurons were the most consistently implicated cell-type across all four psychiatric subgroups and the most affected interneurons and glutamatergic neurons belong to superficial layer 2/3 and 3/4, respectively.

### Most chromosomes are associated with psychiatric cluster disorders

Disease-associated gene-sets can be biased towards specific chromosomes, abnormalities in which can potentially explain the psychiatric disorders within each subgroup. Thus, we looked for chromosomal overrepresentation within the psychiatric subgroups (Fig 4). 15/23 chromosomes showed overrepresentation across the psychiatric cluster, the densest of which were chromosome-5, 8, 12 and 20. Chromosome-5, was overrepresented in disorders associated with addiction (cocaine and alcohol) and autism. Chromosome-8 showed the most diversity and was associated with disorders across three subgroups. Chromosome-12, was overrepresented exclusively in Subgroup-1, representing mood and depressive disorders. Chromosome-20 was overrepresented in subgroup-2 and complex-partial-status-epilepticus in subgroup-3.

**Figure 4:**
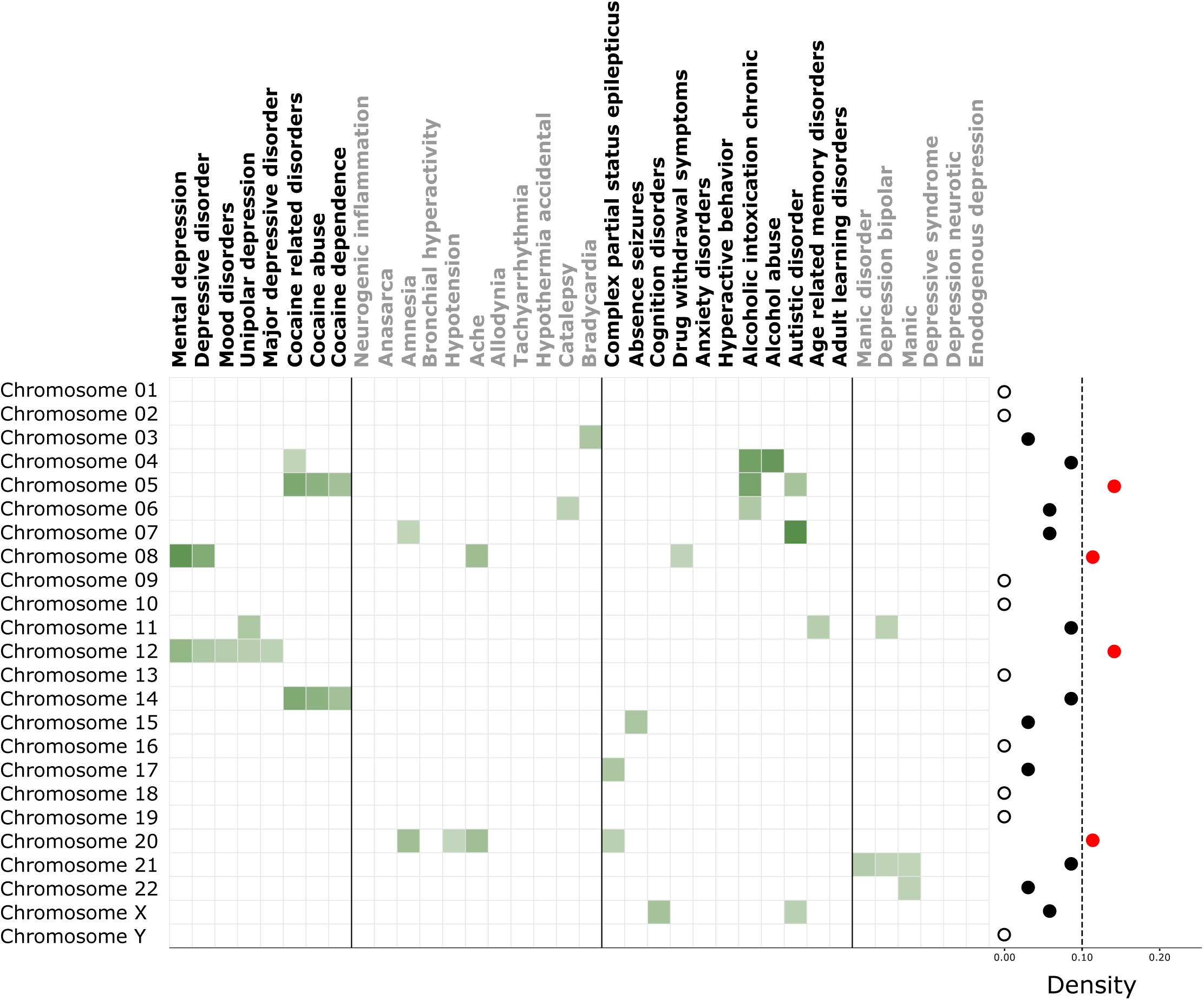
Chromosome overrepresentation analysis of psychiatric cluster disorders. Overrepresentation of different chromosomes across psychiatric cluster disorders (top labels). The filled red dots represent high density chromosomes whereas hollow dots represents chromosomes with zero density.

**Figure 5:**
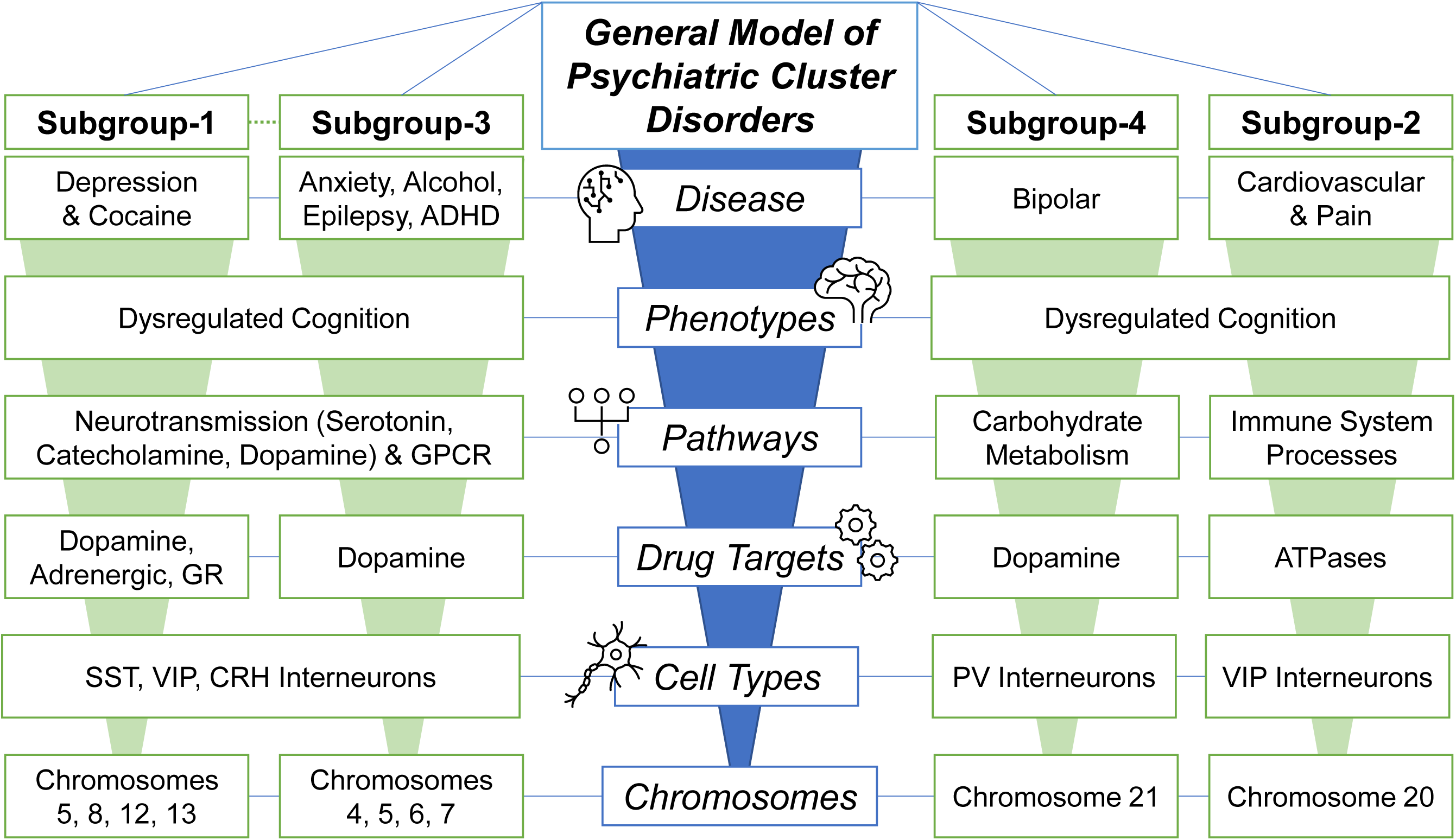
General model of psychiatric cluster disorders. Overall, the present analyses suggest that psychiatric disorders are impacted at physiologic levels of biological complexities, with changes at the chromosome level emanating up to cell-type, drug target, and pathway levels. Collectively these alterations generate various phenotypes, largely related to cognition, which contribute to disease. The present subgroups show similarities and differences across all of these levels, with Subgroup-1 and Subgroup-3 showing the most similarities, and Subgroup-2 showing the most differences.

Within each subgroup, Subgroup-1 showed association with five chromosomes overrepresented across all 8/8 disorders observed here. Interestingly, a split developed within this subgroup, with chromosomes-8 and 12 being more related to depression and chromosomes-5 and 14 being more related to cocaine addiction. Subgroup-2 showed association with five chromosomes across 5/11 diseases observed here, with chromosome-20 being most affected. Subgroup-3 was the most diverse group associated with 9 chromosomes overrepresented across 8/11 disorders observed here. Autism and cognitive disorder, belonging to this subgroup, were the only disorders overrepresented in X-chromosomes.

Subgroup-4 was least diverse with only 3 chromosomes overrepresented across 3/6 disorders observed here. Chromosome 21 showed the highest density in this subgroup associated with bipolar and associated disorders.

### Chromosomal overrepresentation drives the similarity between disorders in psychiatric cluster

Next, we reasoned that, of the four variants –pathways, cell-type, drug-target, and chromosomes— the one which drives the similarity between disorders, should cluster them in agreement with the psychiatry-cluster (Fig 1B, green; Fig S4). Based on rand-index (RI), a measure of similarity between two clusters^27^, we observed moderate but significant similarity between psychiatry cluster and chromosome overrepresentation-based clustering of disorders (RI= 0.70, p-value<3×10^-03^). A weak trend level similarity was also observed between psychiatry cluster and cell-enrichment based clustering (RI= 0.65, p-value<0.07). Finally, among different variants, consistent with the above results, moderated but significant similarity was observed between pathway and MOA based clustering (RI= 0.70, p-value<1×10^-04^). Thus, the similarity between disorders, to some extent, can be explained based on chromosomal instability and cellular correlates of the disorders.

## DISCUSSION

The search for common molecular mechanisms within the wide range of psychiatric disorders is an important future direction. Here, we show that disease specific gene-sets are not only sufficient to segregate diseases based on their intrinsic nature and point similarities between comorbid conditions, but also to move towards a more mechanistic understanding of neural connectivity. Starting with over 750 curated disease-associated gene-sets from DisGeNET, we uncovered three distinct disease clusters (Fig 1A) which shows a fundamental difference between psychiatric/metabolic-type disorders and cancer. Focusing on a psychiatric-cluster branch, we looked for similarity and dissimilarities associated with pathways, drug-targets, cell-types and chromosomes. The most commonly dysregulated process across all disorders involved neurotransmission, neuromodulation and synaptic signaling. Whereas those most unique involved *immune-system-response* and signal-response coupling involving *cell-surface-based-signaling, intracellular-signal-transduction* and downstream response involving *gene-expression* and *metabolic-process*. Independently, the ubiquitous role of neurotransmission was also observed in drug-target based enrichment analysis. In a separate in-silico analysis we showed that the similarity between the disorders is significantly driven by chromosome-based over-representation and to a lesser extent with cell-type based enrichment. This suggests that similarity between disorders disperses in a bottom up fashion i.e. following the subcellular chromosomal abnormality the similarity persists at cellular level where it is affected most by neurotransmission and modulation and disperses at pathway levels possibly through different outside stimulus. Notably, the observed chromosomal-overrepresentation was consistent with large body of literatures on abnormalities of chromosome-5 linked to substance abuse^28,29^, chromosome-8 and 12 linked to lifetime major depression^30,31^ and anxiety disorder^32^ and chromosome-20 linked to epilepsy^33,34^. Further support for similarity at chromosome level comes from genome wide linkage study suggesting shared effects on major psychiatric disorders^4^.

### A general model of psychiatric cluster disorders

Cognition was most affected across all psychiatric-cluster disorders and its known pathophysiological association with neurotransmission can be seen in Figure 2 and further supported by coherent enrichment of dopamine related drug-targets. Within different neurotransmitters, although at different densities, all subgroups showed the highest density of catecholamine— a monoamine group of neurotransmitters. Among other neurotransmitters, serotonin related pathways were densest in the highly correlated subgroups-1 and 2, representing mood and addiction, respectively but not in subgroups-2 and 4, representing neuroinflammation and bipolar disorder, respectively. Overall suggesting that while neurotransmission is a common mechanism between these disorders, the differences may emerge due to different neurotransmitter systems. This raises the question on how things are governed at cell-type level, which as suggested by our results, also drives similarity between the disorders to some extent. Notably, at cell-type level except for ache and bipolar-disorders which showed enrichment of PV positive interneurons, all most all disorders were enriched with dendrite targeting SST and VIP interneurons suggesting that these disorders are largely related to context (all inputs except the one of interest) dependent integration of informational-input to pyramidal neurons, a function largely associated with these neurons^35^. Note that PV neurons, unlike SST and VIP neurons, largely target the axon-initial segments and govern adaptation of output from pyramidal neurons^36^. As such, its enrichment in bipolar disorders aligns with its previously observed similarity with schizophrenia, a disorder associated with abnormal output^2,3^. VIP interneurons by disinhibiting the SST interneurons, also influence the impact of mostly noxious or negative information input^37^. In this regard, their highest enrichment in subgroup-2, suggests their influence on ache, allodynia, tachyarrhythmia– comorbid disorders involving a noxious stimulus of pain. Finally, consistent with the role of corticotropin-releasing hormone in stress, we also observed the enrichment of CRH positive interneurons which are mostly co-expressed with SST or VIP interneurons^15^.

The information input coming to these cell-types are potentially long-ranged as suggested by higher enrichment of monoamine transmitter with origin at distant brain regions or lateral input from different cortical area as suggested by overall enrichment of glutamatergic-transmission. Further support for long distance input comes from the enriched VIP neurons which are influenced by long distance serotonergic and cholinergic afferents^37^. Finally, the enrichment of neuronal markers across all layers also suggests a broad-spectrum behavioural, cognitive and vegetative input from different areas of the brain^38^. However, layer 3/4 receiving thalamocortical input^39^ showed high density for excitatory neurons whereas layer 2/3 responsible for inhibiting layer-1 re-entrant connections^40^ from adjacent areas showed high density for SST interneurons. Overall, the cell and layer-specific enrichment suggests that disorders in the studied psychiatric clusters disorders are influenced by lack of adaptations to input coming from behavioural, cognitive, or vegetative contexts.

### Limitations

This top-down analysis was conducted using unbiased disease associated gene-sets. However, the gene-sets used do not include directionality (upregulation or downregulation). The conclusions we drew for cellular-associates are under the assumptions that the gene-sets used in the study are universal signatures of these disorders regardless of differences in brain region.

