## Supplementary Information for "Similarities and dissimilarities between psychiatric cluster disorders"

**Table of Contents**

**Supplemental Figures:**

- **Figure S1:** Cluster-2 hierarchical dendrogram labeled (corresponds to Fig 1B)
- **Figure S2:** Theme/density correlations between psychiatric cluster subgroups
- **Figure S3:** Drug target frequency across psychiatric cluster
- **Figure S4:** Cluster comparisons and Rand index

**Supplemental Tables (each as separate excel file):**

- **Table S1:** Jaccard matrix for all disease-associated gene sets and corresponding assignments to Cluster-1, Cluster-2, and Cluster-3
- **Table S2:** Detailed results of pathway analysis (corresponds to Fig 2)
- **Table S3:** Detailed results of drug target analysis

**SUPPLEMENTAL FIGURES**

**Figure S1: Cluster-2 hierarchical dendrogram labeled**. Disorders corresponding to red blue and green branch of Cluster-2 dendrogram presented in Fig 1B

**Figure S2:** **Theme-density correlations between psychiatric cluster subgroups**. Correlation coefficient (r) are presented above the diagonal and correlation plots are presented below the diagonal. Each dot represents the theme-density

**Figure S3:** **Drug-target frequency across psychiatric cluster**. Proportional representation of different drug-target in the psychiatric cluster as a whole and within each subgroup. Only top drug-targets are shown.

**Figure S4:** **Comparing hierarchical dendrogram for Rand-index based similarity**. **Top**: comparison of psychiatry cluster dendrogram (corresponds to figure 1B, green) with chromosome and cell-type based dendrogram. **Bottom:** comparison of pathway-based dendrogram with drug-target based dendrogram. Each dashed line (red) shows the dendrogram-cut and corresponding subgroups (black and gray text) of disease labels on the right. In order to generate significance (see methods), for each row (top and bottom) the cluster labels of dendrogram on the left of dashed line (black) was used as reference for resampling based premutation test. Note, only significant comparisons are shown.

**SUPPLEMENTAL TABLES (each as separate excel file)**

**Table S1:** Jaccard matrix for all disease-associated gene sets and corresponding assignments to Cluster-1, Cluster-2, and Cluster-3 (corresponds to Figure 1A)

- **Worksheet-1 [JaccardMatrix (JM)]:** Jaccard matrix for all disease-associated gene sets. Similarity measurements for initial 763 disease-associated gene sets. Values correspond to percent similarity between gene-sets
- **Worksheet-2 [DiseasePerCluster]:** List of all disease-associated gene sets in each cluster, as determined by principle component analysis (corresponds to Figure 1A)
- **Worksheet-3 [JM_Cluster1]:** Jaccard matrix for Cluster-1
- **Worksheet-4 [JM_Cluster2]:** Jaccard matrix for Cluster-2
- **Worksheet-5 [JM_Cluster3]:** Jaccard matrix for Cluster-3

**Table S2:** Detailed results of pathway analysis (corresponds to Figure 2).

- **Worksheet-1 [All_Pathways]:** All unfiltered pathways (row) and corresponding -log10(q-values) per disorder (column) is provided. Column “AM” shows the density of each pathway across all disorders and column “AN” shows the disorder with highest -log10(q-values) value.
- **Worksheet-2 [Filtered_Pathways]:** Pathway filtered for corresponding themes (column “AM” and left labels in Figure 2).
- **Worksheet-3 [ThemeDensity_Index]**: Theme density across all disorders and individual subgroups. The correlation plot matrix for each theme density is shown in figure S2.

**Table S3:** Detailed results of drug target analysis.

- **Worksheet-1 [Drugs-MOA]:** Description (Molecule, MOA, MOA slim, CMap class and target) of all the 132 significantly enriched drugs downloaded from connectivity map data base. Note that MOA slim annotations do not have directions (agonist/antagonist or inhibitor/enhancer) associated with it. As the gene-sets used in the present study do not have directions (up- or down-regulated), we used only MOA slim to describe all the drugs in this study
- **Worksheet-2 [Drugs-DiseaseMOA_all]:** Drug (rows) and corresponding -log10(q-values) per disorder (column) is provided. A summary of the frequency of each drug target is provided in columns AJ and AK.
- **Worksheet-3 [Drugs-DiseaseMOA1]:** Drug-target analysis for Subgroup-1
- **Worksheet-4 [Drugs-DiseaseMOA2]:** Drug-target analysis for Subgroup-2
- **Worksheet-5 [Drugs-DiseaseMOA3]:** Drug-target analysis for Subgroup-3
- **Worksheet-6 [Drugs-DiseaseMOA4]:** Drug-target analysis for Subgroup-4
