## Supplementary figures and images for "Similarities and dissimilarities between psychiatric cluster disorders"

### Figure S1

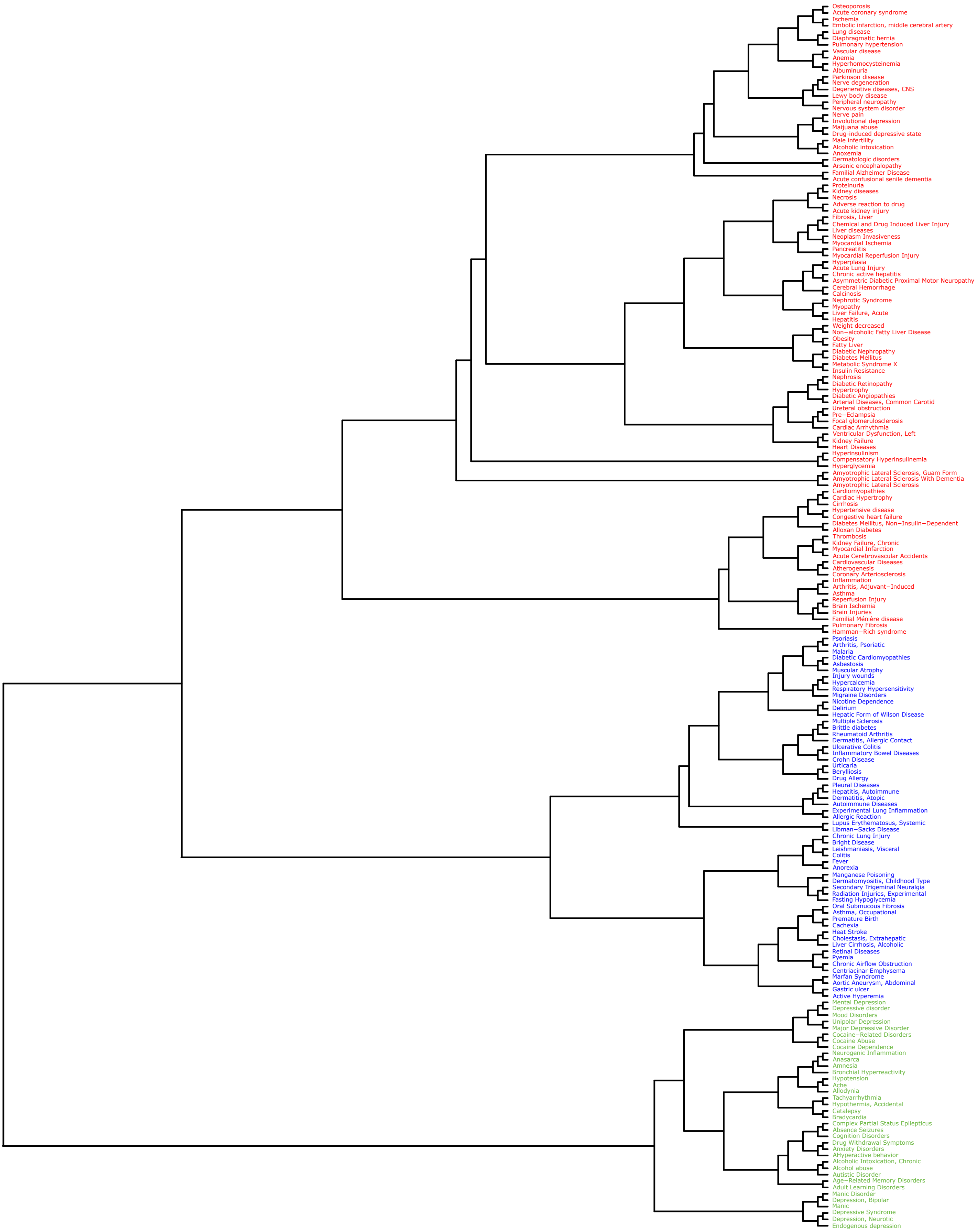

### Figure S2

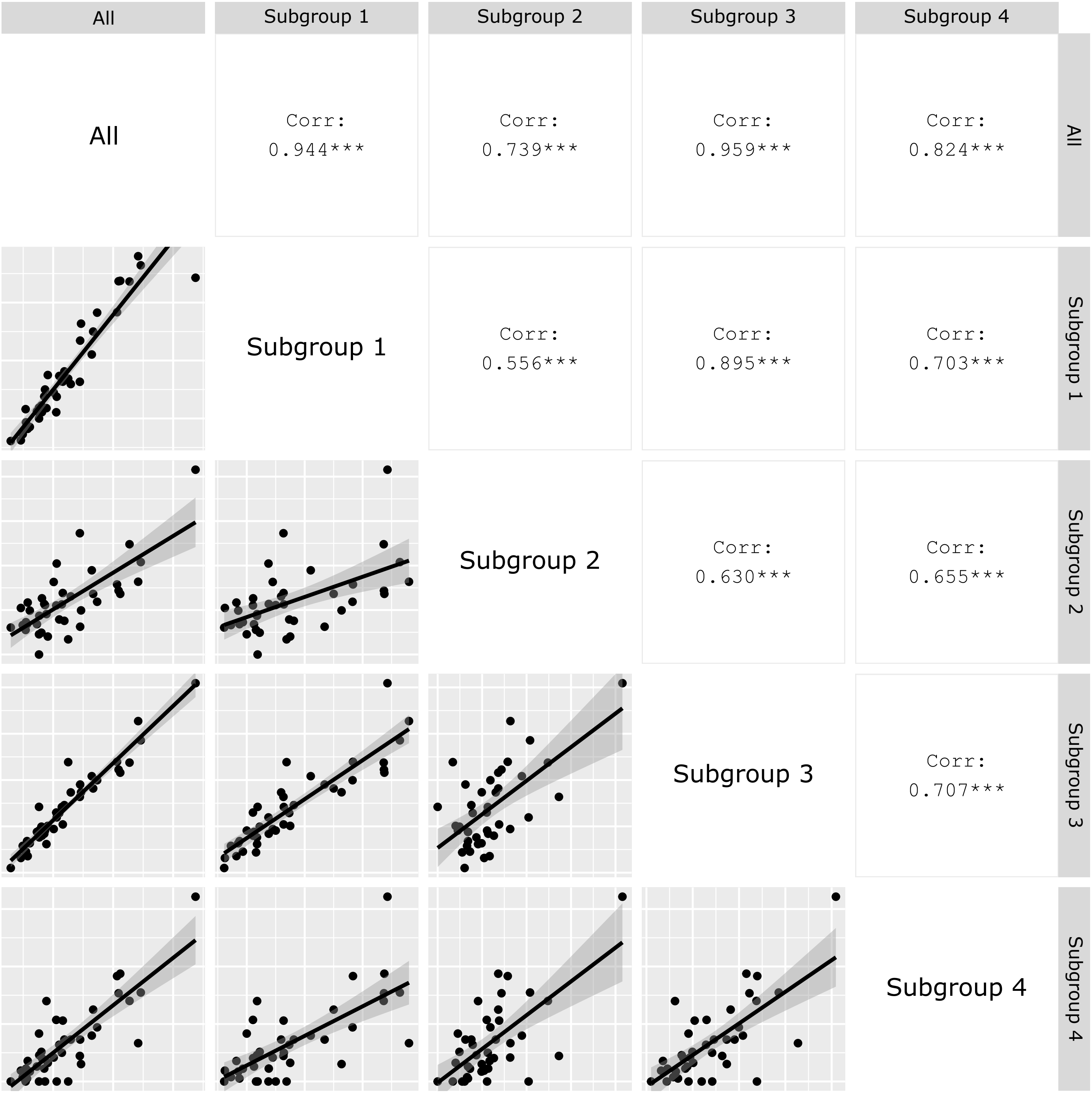

### Figure S3

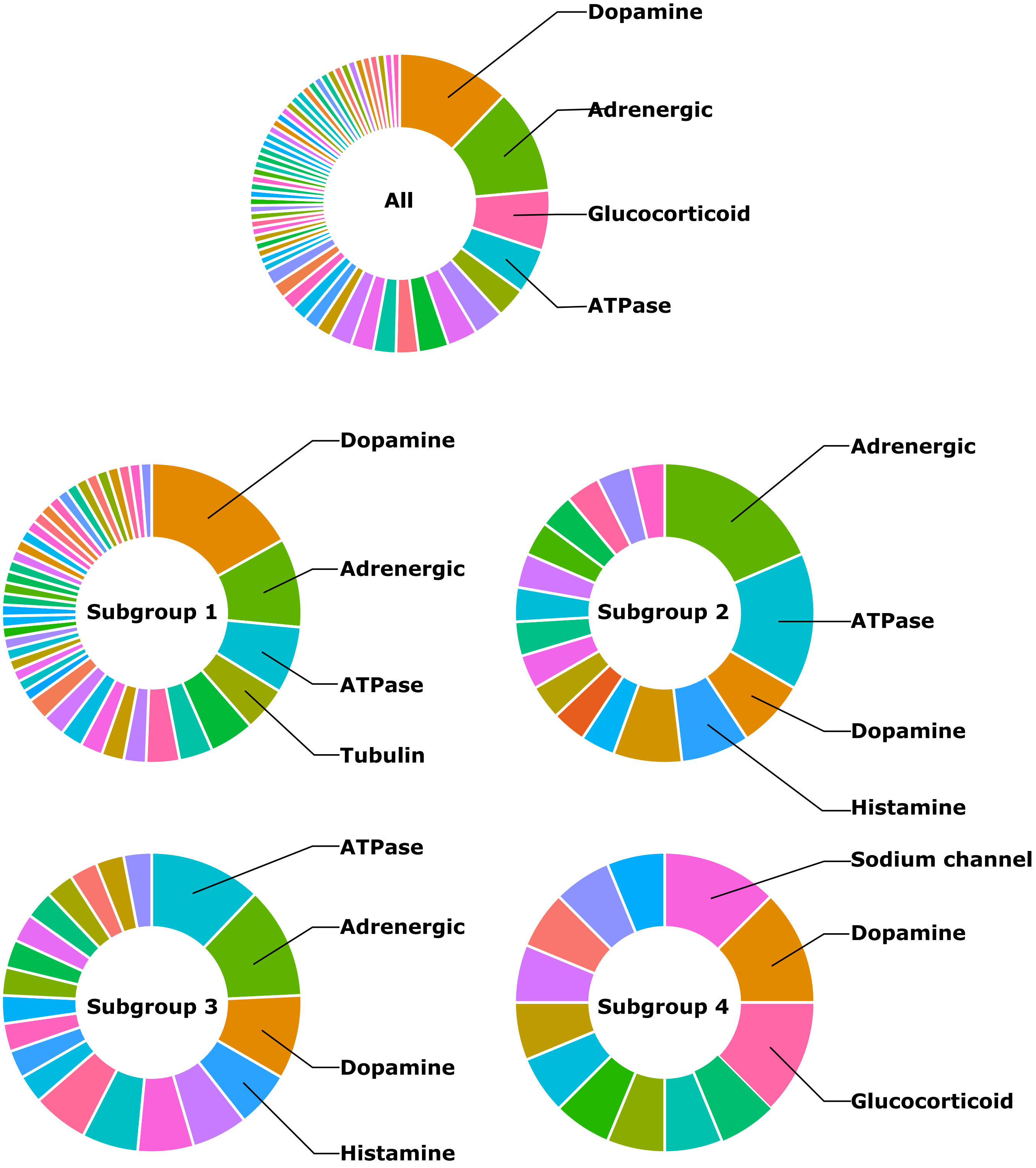

### Figure S4

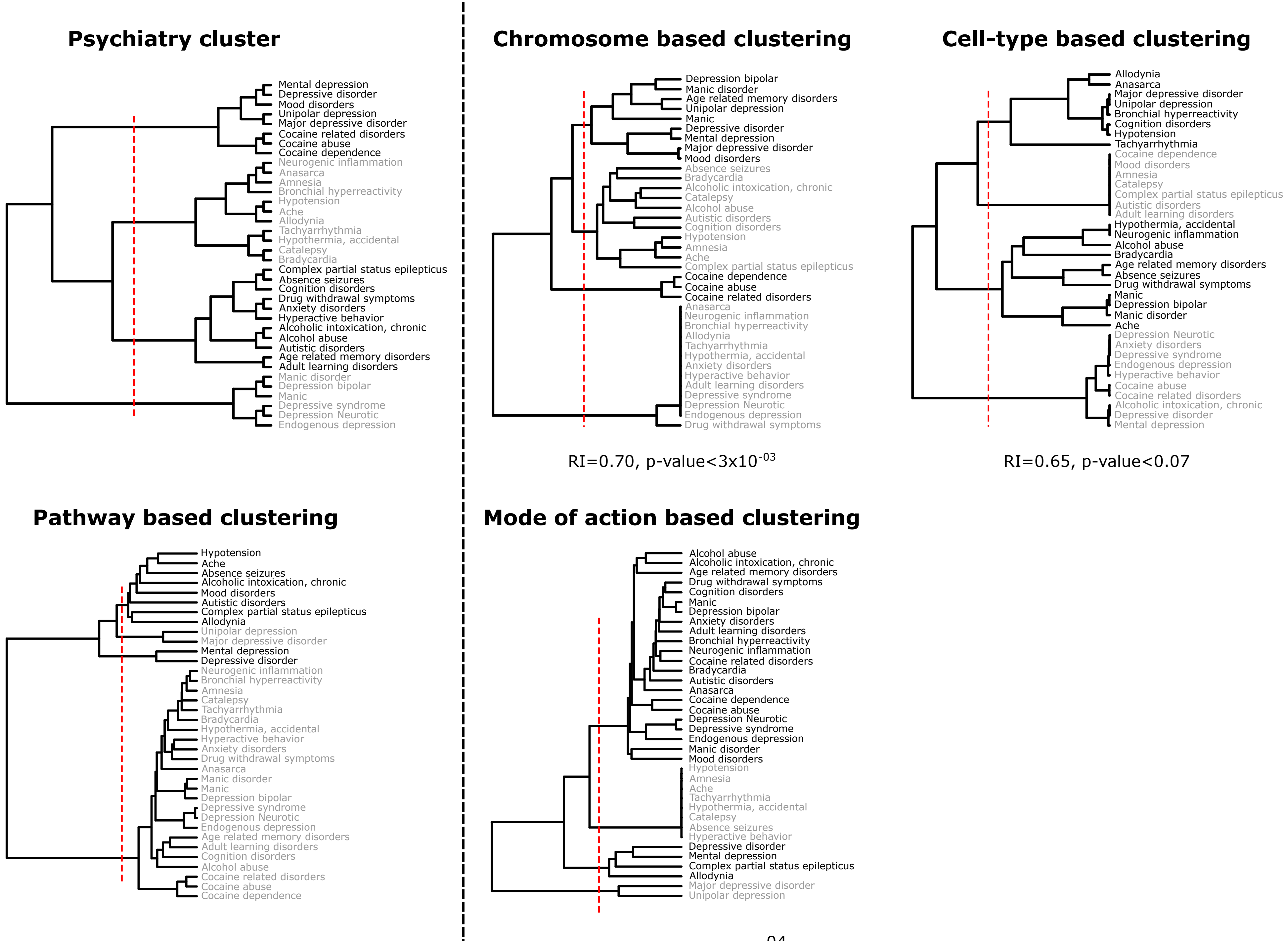
